# Comprehensive identification and characterization of candidate effector proteins in *Puccinia triticina* reveals insights into the wheat leaf rust pathogenesis

**DOI:** 10.64898/2026.04.26.720865

**Authors:** Ankita Shree, Priyanka Kumari, Hasnain Raghib Hassan, Shailendra Kumar Jha, Manish Kumar, Kunal Mukhopadhyay

## Abstract

The biotrophic pathogen *Puccinia triticina* is the causative agent of the most vulnerable foliar disease, namely leaf rust disease of wheat. The pathogen-secreted effectors are essential in modulating fungal virulence and host immune responses. Despite their significance, potential effectors and their underlying mechanisms governing host susceptibility remain elusive. In the present study, we employed an *in silico* approach to identify and characterise effector proteins from the *P. triticina* proteome. Later, performed temporal expression profiling to prioritise effector candidates associated with rust disease. Here, a total of 273 high-confidence effector candidates were identified and analysed their physicochemical properties, domains, motifs, and functional annotations, to assess their conservation and dynamics. Although most of the effectors were uncharacterised, the conserved motif virulence-associated [YFW]xC was notably enriched in the effector repertoire. Comparative PHI-base annotation highlighted similarities with known fungal virulence factors involved in host susceptibility. Effectors harbouring CAZyme activity indicate involvement in host cell wall modification. Promoter analysis identified multiple stress- and defence-related transcription factor binding sites, suggesting regulated expression during infection. Transcriptome analysis revealed that 20 effector genes were significantly upregulated during *P. triticina* infection. qRT-PCR validated the expression of 4 highly induced effector transcripts following *P. triticina* infection in susceptible wheat variety. Specifically, two of these candidates demonstrated biphasic expression pattern that aligns contrasting PTI- and ETI-mediated defense mechanisms critical for sustained virulence. Overall, this study provides a comprehensive framework for identifying functionally relevant *P. triticina* effectors and offers insight for future effector-target studies and effector-based leaf rust management strategies.

## Introduction

Plants, being sessile organisms, are exposed to numerous abiotic and biotic challenges. Biotic pathogens such as viruses, bacteria, fungi, nematodes, and aphids reside in close proximity to the host plant and pose a significant threat to crop health and productivity. Among these biotic pathogens, fungal phytopathogens account for approximately 70% of plant diseases, thereby concurrently affecting the economic growth of the country (Mahadevakumar and Sridhar, 2021). Phytopathogenic fungi invade their host cells via the growing hyphal tip, a process that compromises fungal virulence (Sinha et al., 2021). Nevertheless, the multilayered defense systems of plant species enable their persistence and survival against all odds (Jones and Dangl, 2006). Plants detect conserved pathogen-derived molecules, known as pathogen-associated molecular patterns (PAMPs), such as flagellin Flg22 and chitosan, through their cell-surface-resident pattern recognition receptors (PRRs). This recognition activates the plant’s innate immune response, including the production of reactive oxygen species (ROS), callose deposition, initiation of the miRNA pathway, downstream signalling cascades, and the induction of defence-related gene expression. Collectively, these events constitute the first line of broad-spectrum defence against invading pathogens and are referred to as PAMPs-triggered immunity (PTI) (Jones and Dangl, 2006; Jonge R et al., 2006; Navarro et al., 2008).

Although PTI is prevalent in plants, fungal pathogens evade or suppress these sophisticated surveillance systems and downstream defences by secreting an arsenal of effector proteins that promote infection. Secreted effectors can enter various cellular compartments of host cells, including the cytoplasm, nucleus, and chloroplast (Jones and Dangl, 2006; Bigeard et al., 2015). In these compartments, they interact with their respective host targets or act as molecular decoys, thereby disrupting the normal function of these targets. These interactions are crucial for manipulating the host’s basal defence response and facilitating the successful colonization of the pathogen within the host. For instance, Pst12806, an effector derived from the stripe rust pathogen, is translocated into the chloroplast and undermines basal immunity (Xu et al., 2019). Highly specialized Nucleotide-binding Leucine-rich Repeat (NLR) proteins recognise some of these host-translocated effectors and trigger a robust immune response, including hypersensitive response (HR) (Prasad et al., Jones et al., 2016). Notably, effectors undergo rapid evolution, which significantly contributes to the swift evolution of virulence among adapting fungal pathogens (Sahu et al., 2024).

However, for many pathogens, the underlying signalling networks targeted by effectors and the functional significance of these interactions remain poorly understood. Chloroplasts integrate various environmental stimuli and play a central role in plant defence. In addition to photosynthesis, chloroplasts produce ROS for antimicrobial activity. However, enhanced immunity often compromises crop yield (Huot et al., 2014). For biotic pathogens that rely on living hosts for nutrient acquisition, it is important to create a favourable environment that sustains both growth and yield. Given their importance, chloroplast proteins have emerged as a key target for manipulation by pathogens (de Torres et al., 2015).

Wheat (*Triticum aestivum* L.) is a widely grown crop that provides protein and calories to more than 50% of the world’s population (Kiran et al., 2016). However, global productivity of wheat is majorly threatened by the devastating biotrophic fungi *Puccinia triticina*, the causal agent of macrocyclic foliar rust disease. The disease incidence is reported in all the wheat-producing countries, and it is the most prevalent wheat disease in India (Bhardwaj et al., 2010). The availability of genomics and transcriptomics strategies has improved our understanding of the roles of differentially expressed genes in host-pathogen interactions across multiple stages of *P. triticina* infection. Approximately 80 Leaf rust resistance (*Lr*) genes have been catalogued in wheat, conferring resistance against *P. triticina*. Notably, multiple pathotypes remain avirulent towards Indian leaf rust differentials carrying *Lr*9, *Lr*10, *Lr*19, *Lr*24, and *Lr*28 (Prasad et al., 2017; Sharma et al., 2018). Understanding the molecular crosstalk between pathogen effectors and host targets is key to decoding mechanisms of pathogenesis and developing disease-resistant cultivars.

Thus far, systematic studies on *P. triticina* secreted effectors and their role in virulence are in their infancy. Although numerous effector candidates have been predicted in *P. triticina*, many remain uncharacterized, and their host targets are unknown. Studies have identified effector candidates in *P. triticina* using transcriptomic and motif-based approaches (Zhao et al., 2020; Prasad et al., 2023). These strategies may overlook effectors with dynamic expression patterns or lacking conserved sequence features. To overcome these limitations, we employed a comprehensive secretome analysis that integrates key effector-associated characteristics for complete effector prediction, followed by RNA-seq data analysis to identify candidate virulent effectors of *P. triticina*. Overall, this comprehensive study aims to provide essential insights into wheat-rust interactions, with potential implications for crop improvement and global food security.

## Materials method

### Genome-wide prediction of P*. triticina* Effector protein

Proteome data of *P. triticina* was retrieved from Ensembl Fungi database (https://fungi.ensem bl.org). *Puccinia triticina* GCA000151525v2 was used for effector identification and further *in silico* analysis involving online available machine learning tools. For secretome analysis and to identify potential effector candidates, initially secretory proteins were sorted from the total fungal protein. Therefore, 15,685 *P. triticina* proteins were subjected to SignalP 5.0 to determine proteins with N-terminal signal peptides. Then, secretory proteins were screened for the absence or presence of a transmembrane domain. TMHMM 2.0b predicted proteins with 0 or 1 transmembrane domains. Later, Phobius v1.01 was used to refine signal peptide and TM predictions. Protein with Signal peptide and absence of transmembrane domain were used as input in WoLFPSORT v0.2 to predict extracellular localization. Further, TargetP 2.0 filtered proteins with mitochondrial or chloroplast transit peptides and PROSITE (PS00014) excluded proteins with ER retention signals. To obtain *P. triticina* secretory protein, PredGPI was used to eliminate proteins with glycosylphosphatidylinositol (GPI) anchors. Finally, proteins were screened for size (≤400 amino acids), cysteine content (≥3%), and analyzed using EffectorP 3.0 to filter *P. triticina* effector protein from the subset of secretory proteins.

### Phylogenetic analysis of effector proteins, Gene Structure Organization, and Conserved Motif Identification

Multiple sequence alignment was carried out using the Muscle tool with default parameters. Subsequently, phylogenetic analysis was inferred using neighbor-joining method in Mega7. The robustness of the tree topology was evaluated by bootstrap values with 1000 iterations (Kumar et al., 2016).

The genomic and corresponding coding sequences (CDS) of effectors were retrieved from the reference genome annotation file (GTF/GFF format) available at the Ensembl fungi database. The exon-intron boundaries were visualised using TBtools. The number, length, and distribution of exons and introns were compared among the effector genes. In this study, conserved effector motifs like RxLR, [YFW]xC, and others were identified using a regular expression (RegEx)-based approach. The pattern matching was performed using custom Shell scripts, in which regular expressions were applied to identify motif occurrences within protein sequences.

### Physicochemical analysis and subcellular localization prediction of effector protein in host

The comparative physicochemical properties of effector proteins were analyzed and compared with those of the total secretory proteins. Parameters including sequence length, amino acid composition, molecular weight (MW), isoelectric point (pI), instability index, aliphatic index, and grand average of hydropathicity (GRAVY) were elucidated using the ProtParam tool available on the expert protein analysis system (ExPASy) server (http://www.expasy.org). Additionally, the hydrophilic or hydrophobic nature of proteins was evaluated based on GRAVY values. Furthermore, cysteine residues in effectors and secretory protein were computed through custom commands in Linux.

Host localization of effector proteins was determined from the online available tool LOCALIZER (https://localizer.csiro.au/). Signal peptides from effectors are cleaved off before delivery of the effectors to the host. Therefore, the protein sequence without SP was submitted as query for prediction.

### Promoter Scanning

The promoter sequence of 1000 bp upstream of the translation start site of each respective *P. triticina* gene was retrieved from the Ensembl Fungi database. Initially, the transcription factor binding motifs were identified using position frequency matrices obtained from the JASPAR 2026 CORE fungal database. Followed by Motif scan using the FIMO (Find Individual Motif Occurrences) program available at MEME Suite (v5.5.8). The fungal non-redundant JASPAR motif collection was used as input, and promoter sequences were scanned using default settings. Significant motif occurrences were filtered based on a threshold of *p* < 1e-4.

Motif distribution along promoter regions was illustrated using the “Visualize Gene Structure (from GFF/GTF file)” module in TBtools (Chen et al., 2020). Transcription factors were further categorized into functional classes based on their DNA-binding domains, including bHLH, C2H2 zinc finger, bZIP, MADS-box, C6 zinc cluster, HMG, homeodomain, forkhead/winged helix, and others.

### Structure Analysis and functional annotation of effector proteins

The structure of highly expressed effectors was predicted using the I-TASSER server. Protein sequences were submitted in FASTA format to the I-TASSER server. The server predicted three-dimensional structures from the template protein in PDB database. The selected 3D models were visualised using PyMOL molecular visualization tool. Structural features such as α-helices, β-sheets, and surface properties were examined to understand potential functional regions and interaction interfaces.

Functional annotation of effectors was carried out using InterProScan v5.47 82.0 to obtain Gene Ontology (GO) terms and Pfam domain assignments. Thereafter, the results were integrated with OmicsBox platform, to assign GO terms based on sequence homology and domain architecture. CAZymes were identified using automate web tool dbCAN3 (https://pro.unl.edu/dbCAN2).

### *In silico* expression analysis of secreted protein from transcriptome data

RNA-seq data of *P. triticina* plant–pathogen interactions analaysed here were obtained from NCBI Bioproject, with corresponding accession numbers (PRJNA328385 and PRJNA182354). Clean reads were aligned to fungal genomes using HISAT2. Gene expression was calculated using subreads, and fold changes for differentially expressed genes (DEGs) were determined. Further, the list of protein candidates of this study was screened from DEGs. Heatmaps were generated in TBtools using a normal count file (Chen et al., 2020).

### Wheat plant growth and rust inoculation

Wheat NIL plant HD2329 +Lr28 (seedling resistant to leaf rust) and HD2329 (seedling susceptible to leaf rust) were grown until the two-leaflet stage in pots containing soil and farmyard manure (7:1 ratio) within a controlled environment in the greenhouse at IARI, PUSA campus, New Delhi. For infection, urediniospores of *P. triticina* (Pathotype 77-5) were resuspended in sterile water at a concentration of 20 urediniospores/µl. 0.75% Tween 20 was added as a surfactant to the urediniospore solution. To observe rust symptoms, the plants were challenged with urediniospores via spraying. Plants sprayed with sterile water were used as the control plant. To enhance rust disease symptoms, plants were maintained at 95% humidity for 48hr in dark. Afterwards, plants were transferred to greenhouse conditions maintained at: light period 16h D/N, humidity 80%, and temperature 22°C. Leaf samples were collected at 0, 12, 24, 48, 72, and 168 hpi and stored at -80 °C for further use.

### RNA extraction and Relative expression of key effector genes

To evaluate the expression patterns of candidate effector genes potentially involved in *P. triticina* infection, wheat near-isogenic lines HD2329 + Lr28 (resistant) and HD2329 (susceptible) were used in this study. Both resistant and susceptible *P. triticina*, as well as mock-inoculated leaves, were collected at 0, 24, 48, 96, and 168 hpi. RNA was isolated from both infected and uninfected leaves using RNA Isolated Kit (Qiagen), according to the user manual. Reverse transcription was carried out using the First-Strand cDNA Synthesis kit (ThermoScientific), following the manufacturer’s instructions. Expression primers for respective effector genes were designed and *P. triticina* elongation factor was used as internal control. The qRT-PCR experiments were performed with three independent cDNAs, including three technical replicates. Relative fold change was computed using the 2^-ΔΔCT^ method (Paolacci et al., 2009).

## Results

### *In silico* identification and Physicochemical Characterization of *P. triticina* effector proteins

In this study, a conventional bioinformatics pipeline was employed to predict secretory proteins potentially acting as effector proteins in the pathogenic fungi *P. triticina* **(Figure 1A)**. A total of 15685 unique proteins encoded by *the P. triticina* genome, were evaluated for secretory features, including signal peptide, extracellular localization, and absence of transmembrane domain and glycosyl phosphatidyl inositol anchors. Out of the total sequences, **1520** were predicted with high-potential signal peptides, comprising **9.69**% of the *P. triticina* genome. Thereafter, secretory proteins were scrutinised for effector characteristics, including amino acid length ≤ 400 and a cysteine residue content of ≥ 4%, followed by effector prediction using EffectorP. From the initial pool of 1520 secretory proteins, subsequent effector prediction analysis identified 273 as probable *P. triticina* effector proteins, provisionally designated as PtEPs, as they attain the criteria delineated by EffectorP. Overall, PtEPs constitute 19.96% of the total secretome and account for 1.74% of the total *P. triticina* genome. Among the 273 PtEPs, 135 were predicted to function as apoplastic effectors, while 57 were classified as cytoplasmic effectors. Additionally, 81 proteins exhibited features of both categories **(Figure 1B)**. We also investigated the characteristic features of effector proteins, including their physicochemical properties and cysteine content, relative to those of total secretory proteins. Based on the analysis of amino acid sequences, PtEPs exhibited typical cysteine richness and shorter sequence lengths compared to secretory proteins, consistent with the known properties of effector proteins. Precisely, the sequence length of effector protein ranged from 47 to 399 amino acids, and the number of cysteine residues varied from 3 to 22. On the contrary, secretory proteins displayed much broader distribution with sequence length ranging from 43 to 2437 amino acids and cysteine content ranging from 0 to 70 **(Figure 1C)**. Molecular weight distribution of effector and secretory proteins was also evaluated. The molecular weight of effector protein was concentrated in between 5.09 to 43.82 kD with an average weight of 17.33 kD, while for secretory proteins the molecular weight broadly varied from 4.44 kD to 272.07 kD **(Figure 1C)**. Interestingly, no significant differences were observed in properties such as isoelectric point, Instability index, Aromaticity, and GRAVY between secretory and effector proteins **(Supplementary Figure 1)**.

**Figure 1.**
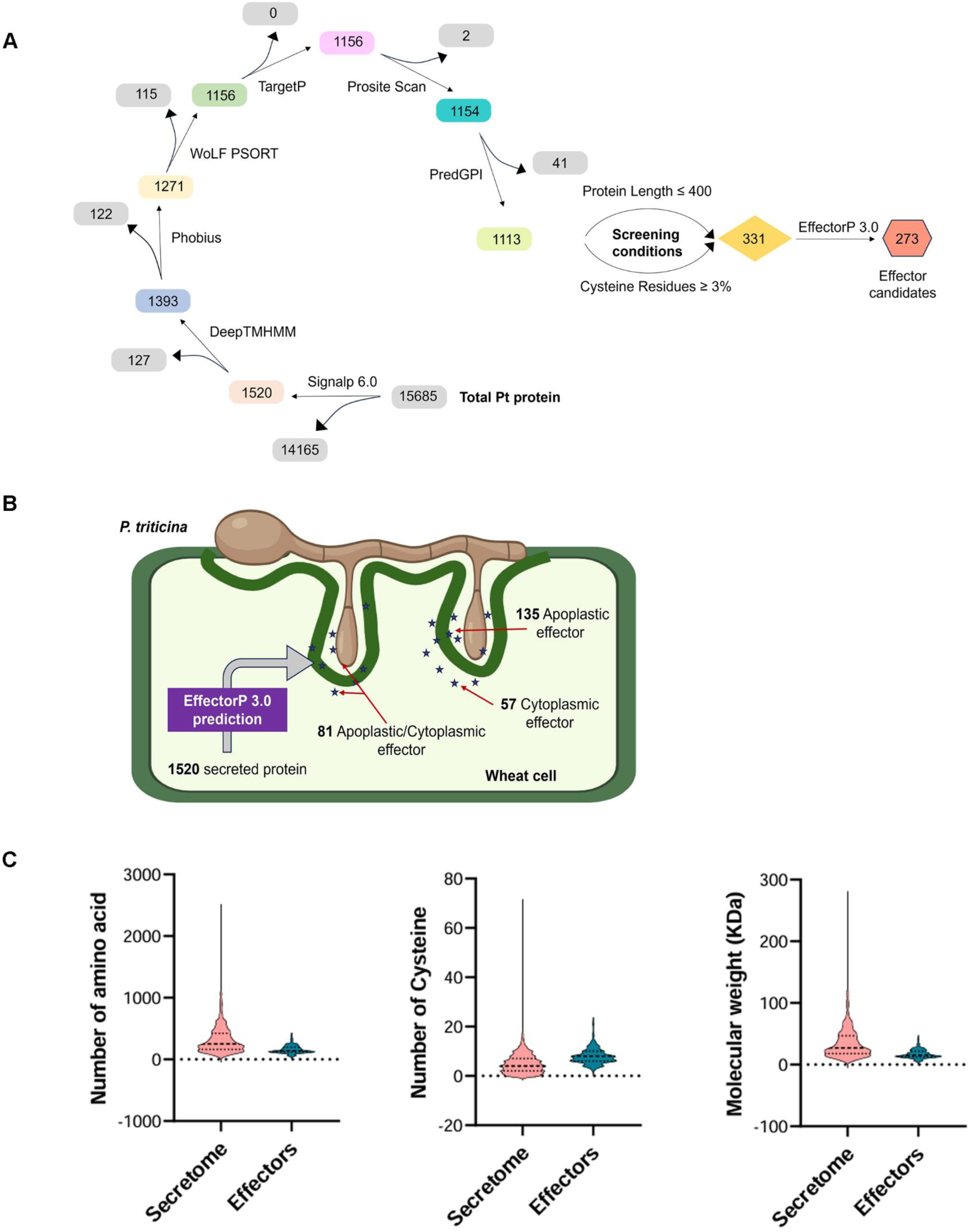
Effector prediction pipeline and sequence feature analysis of identified *P. triticina* effector proteins: (A) Bioinformatics workflow for predicting and analyzing effector proteins in the *P. triticina* proteome. (B) Illustration depicting *P. triticina* penetrating a wheat cell and localization of multiple effectors. (C:i-iii) Comparison of the number of amino acid residues, cysteine residues, and molecular weight distribution in predicted secretory (SP) and effector (EP) proteins.

To gain insight into the evolutionary history of PtEPs phylogenetic analysis was performed using full-length protein sequences. The MUSCLE tool was used to align the protein sequences, and the aligned sequences were used to construct an unrooted phylogenetic tree using Neighbor-joining (NJ) method. The phylogenetic tree categorized all 273 effector proteins in **123** clades, This suggests that effectors within the same cluster probably share similar functions (**Supplementary Figure 2)**. Overall, this robust approach, based on secretory and effector features, including amino acid length and cysteine content, enables the reliable identification of candidate effector proteins from total *P. triticina* secretome.

### A comprehensive understanding of PtEPs characterization with functional annotation

To explore the potential functions of PtEPs, we performed a detailed analysis of sequence features, such as motifs and domains. Screening for known effector-associated motifs revealed that several PtEPs contain conserved signature motifs previously identified in effector proteins. Notably, the [YFW]xC motif appeared in the majority of effector subsets, approximately 57.50% **(Figure 2A)**. The abundance of the [YFW]xC motif in PtEPs correlates with its frequent presence in fungal effectors, suggesting a fungal strategy to evade host defences and to facilitate effector-mediated virulence and host adaptation. Additionally, we identified the RxLx[EDQ] and [LI]xAR motifs in 7.32% and 6.59% of PtEPs, respectively, along with the 2.19% occurrence of the oomycete-specific RxLR motif in a considerable number of instances **(Figure 2A)**. Furthermore, numerous effector proteins possessed two or more motifs concurrently **(Figure 2B)**. To broaden motif identification, conserved motifs were analysed across the entire set of PtEPs. A total of 10 consensus motifs were identified using the MEME programme. Among the analysed sequences, only 44 PtEPs exhibited conserved sequences. Of these, Motif 9 was highly enriched, followed by Motif 3, while Motifs 4, 5, and 8 were the least prevalent **(Supplementary Figure 3)**.

**Figure 2.**
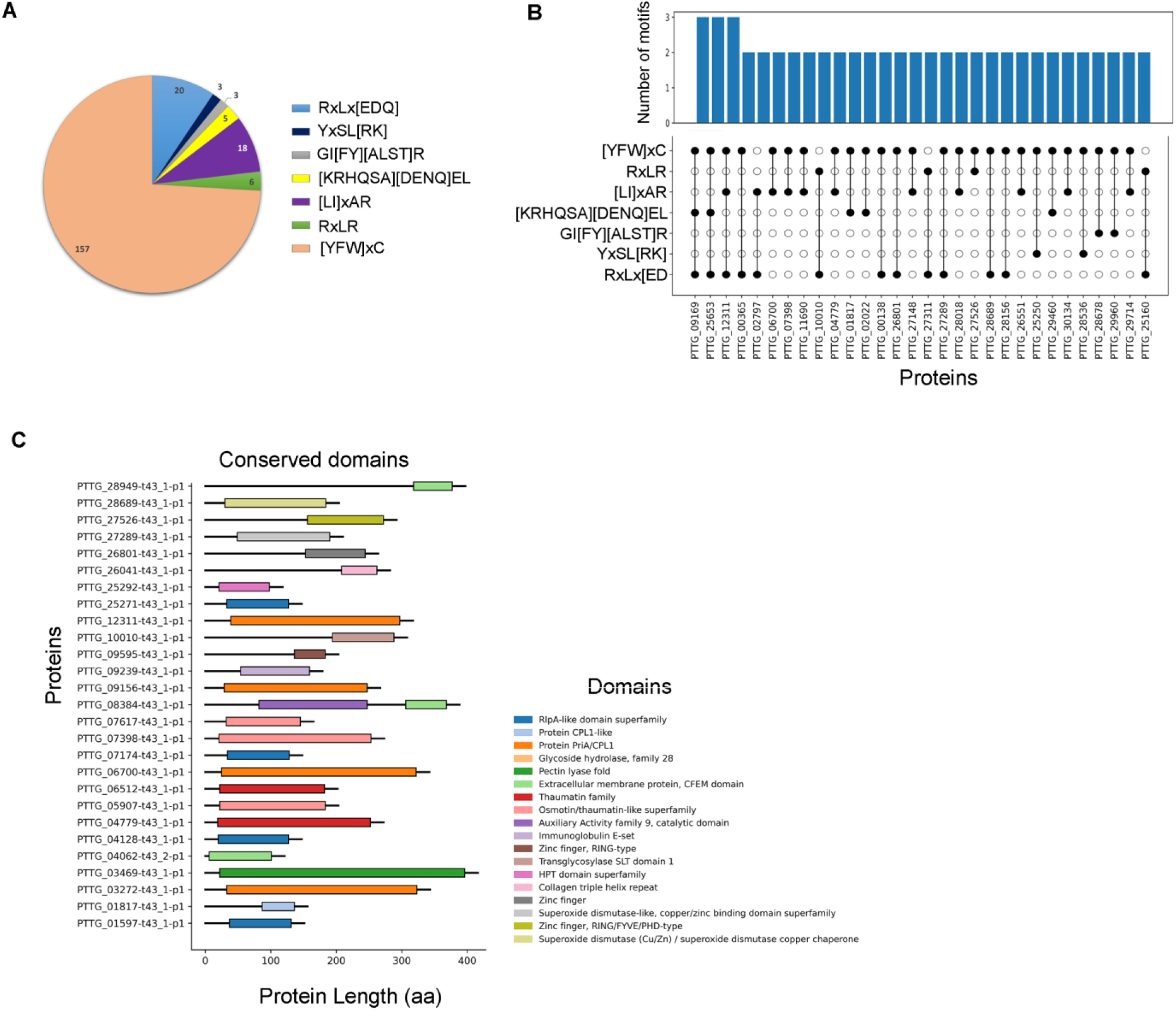
Distribution of motifs and domain architecture of the predicted effector proteins. (A) Distribution and prevalence of conserved motifs within the predicted effector repertoire. (B) A stacked chart showing the presence of two or more motifs within individual effector proteins. The upper panel depicts the quantity of motifs that co-occur, while the lower panel delineates the types of co-existing motifs. (C) Schematic depiction of domain architectures in selected secreted and effector proteins. The scale represents the full-length protein sequence. Coloured boxes depict the position and range of the respective domain(s). Each domain is represented by a specific colour, as described on the right.

To expand the understanding of the functional potential of PtEPs, a conserved domain analysis was performed utilising publicly available databases. The effector sequences were annotated through an extensive array of protein databases, including PANTHER, Pfam, PROSITE, KEGG, and CAZy **(Figure 2C)**. Overall, **1234** effectors displayed sequence annotation. **28** effectors were annotated by PANTHER, **27** by Pfam, and **17** by PROSITE_PROFILES. Among the analysed sequences, conserved domains corresponding to known protein families were identified in only 32 PtEPs via the NCBI Conserved Domain Database (CDD), which accounts for 10.6% of the total PtEPs **(Figure 2D)**. Here, the domain architecture revealed that domains were highly diverse and variably distributed across different proteins. Even the similar domains were located at different positions within the protein sequences. Some of the proteins contained more than one domain as well. These results indicate that the majority of effectors in *P. triticina* are novel or harbour uncharacterized or undetectable conserved domains. Sequence feature motifs consistent with domain organization support both the functional diversity and complexity of the predicted effector repertoire.

To better understand the functional features of 273 PtEPs, we performed Gene Ontology (GO) annotation. This analysis showed that 40 proteins, accounting for about 14.65% of all PtEPs, were annotated with various GO terms. Specifically, 22 terms could be assigned linked to key biological, cellular, and metabolic processes, providing valuable insights into their roles **(Figure 3A**). In the KEGG database, we found that 84 proteins were functionally annotated, which accounts for 30.7% of the total PtEPs **(Figure 3B)**. The results showed that many of these proteins were linked to a variety of important pathways, such as metabolism, genetic information processing, environmental information processing, and cellular processes. Notably, 40 proteins were associated with carbohydrate and amino acid metabolic pathways, suggesting their potential involvement in modulating host metabolic processes. Collectively, these findings strongly suggest that PtEPs function in host interactions, immune modulation, and diverse physiological functions essential for pathogenicity.

**Figure 3.**
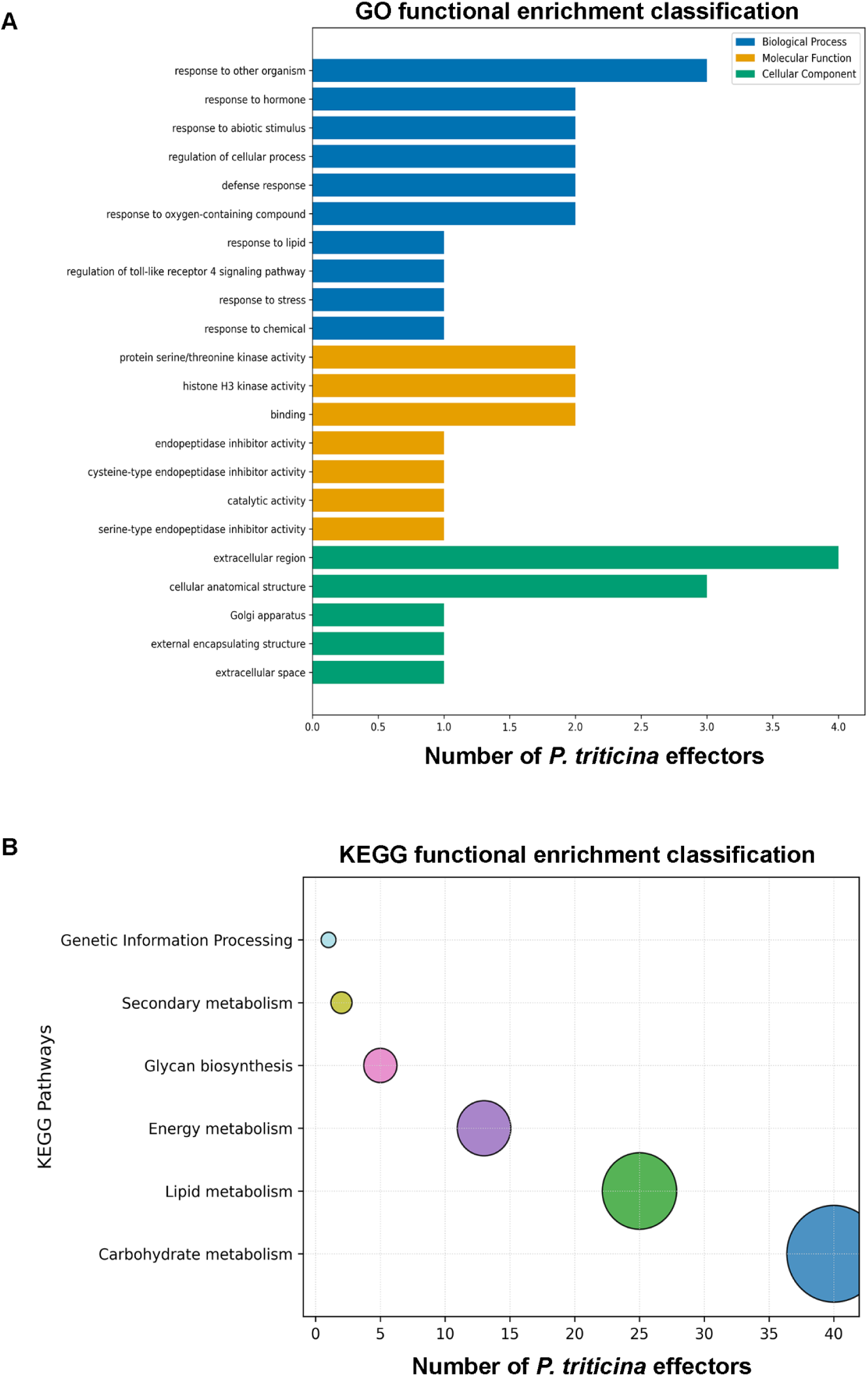
Functional features of the predicted effectors. (A) Bar plot showing enriched Gene Ontology (GO) terms grouped by associated protein domains (B) KEGG annotation for identified effectors. KEGG annotation shows enrichment in carbohydrate metabolism in PtEPs.

Fungal pathogens modify host cell wall components to facilitate pathogen entry and nutrient acquisition. Therefore, we investigated potential effector proteins that could function as carbohydrate-active enzymes (CAZymes) and can degrade or modify host cell components. We performed a CAZyme analysis using the online tool dbCAN2. Here, we identified a total of 9 PtEPs that may have CAZyme functions. Interestingly, 7 of the 9 were glycoside hydrolases (GHs), followed by auxiliary activity enzymes (AA9) **(Table 1)**. Overall, finding CAZymes among these predicted effectors suggests they could be key players in helping the pathogen evade host defences by altering structural barriers and host-derived carbohydrates.

**Table 1:** Identification of Carbohydrate-Active Enzymes in predicted *P. triticina* effector proteins using dbCAN2.

| S. No. | Protein ID | Different Tools |  |  |
| --- | --- | --- | --- | --- |
|  |  | HMMER | DIAMOND | dbCAN_sub |
| 1 | PTTG_03469 | GH28(65-384) | GH28 | GH28_e319 |
| 2 | PTTG_04779 | GH152(26-252) | GH152 | GH152_e44 |
| 3 | PTTG_05907 | GH152(28-169) | GH152 | GH152_e32 |
| 4 | PTTG_06512 | GH152(27-167) | GH152 | GH152_e32 |
| 5 | PTTG_07398 | GH152(27-253) | GH152 | GH152_e44 |
| 6 | PTTG_08384 | AA9(14-247) | AA9 | AA9_e81 |
| 7 | PTTG_28949 | AA9(11-247) | AA9 | AA9_e81 |
| 8 | PTTG_10010 | N | GH23 | GH23_e1371 |
| 9 | PTTG_25283 | N | GH16_1 | N |

To identify experimentally validated effector proteins from other pathogens homologous to *P. triticina* effectors, we analysed the Pathogen-Host Interaction (PHI) database. Nearly 16 predicted effector proteins, representing 5.86% of the total PtEPs, were functionally annotated in the PHI database **(Table 2)**. Among the annotated protein sequences, the majority of the predicted effector proteins showed functional similarity to the “virulence regulators” in *Cryptococcus neoformans*, followed by “oxidative stress response determinant” in *Puccinia striiformis*. This alignment of PtEPs with sequences from different pathogens in the PHI database offers important insight into their possible evolutionary relationships and functions in plant-pathogen interactions. However, fewer significant similarity to characterized virulence proteins in PHI-base, suggests that the majority of predicted *P. triticina* effectors are novel or highly divergent.

**Table 2:** *P. triticina* effector proteins exhibiting sequence similarity to functionally annotated known effector proteins in PHI BLAST.

| S. No. | Sequence ID | PHI base ID | Pathogen | Gene / Product | Identity (%) | E value | Function |
| --- | --- | --- | --- | --- | --- | --- | --- |
| 1 | PTTG_28659 | PHIG:168 | *Puccinia striiformis* | Copper/zinc superoxide dismutase | 37.824 | 2.39E-27 | Oxidative stress response |
| 2 | PTTG_28689 | PHIG:168 | *Puccinia striiformis* | Copper/zinc superoxide dismutase | 32.461 | 4.89E-25 | Oxidative stress response |
| 3 | PTTG_27289 | PHIG:168 | *Puccinia striiformis* | Copper/zinc superoxide dismutase | 37.762 | 1.91E-23 | Oxidative stress response |
| 4 | PTTG_01597 | PHIG:5241 | *Cercospora beticola* | RlpA-like protein double-psi beta-barrel domain-containing protein | 44.248 | 6.33E-22 | Cell wall remodeling |
| 5 | PTTG_07174 | PHIG:5241 | *Rhizoctonia solani* | RlpA-like protein double-psi beta-barrel domain-containing protein | 41.228 | 3.55E-20 | Cell wall remodeling |
| 6 | PTTG_04128 | PHIG:5147 | *Ustilago maydis* | RlpA-like protein double-psi beta-barrel domain-containing protein | 40.196 | 8.10E-19 | Cell wall remodeling |
| 7 | PTTG_25271 | PHIG:5241 | *Cercospora beticola* | RlpA-like protein double-psi beta-barrel domain-containing protein | 42.574 | 4.30E-17 | Cell wall remodeling |
| 8 | PTTG_06700 | PHIG:3617 | *Cryptococcus neoformans* | Protein CPL1 | 39.394 | 4.80E-16 | Signal transduction / virulence regulator |
| 9 | PTTG_12311 | PHIG:3617 | *Cryptococcus neoformans* | Protein CPL1 | 33.898 | 3.38E-15 | Signal transduction / virulence regulator |
| 10 | PTTG_03469 | PHIG:3069 | *Bipolaris zeicola* | Exopolygalacturonase | 25.714 | 7.90E-14 | Cell wall degradation enzyme |
| 11 | PTTG_03272 | PHIG:3617 | *Cryptococcus neoformans* | Protein CPL1 | 36.364 | 9.52E-14 | Signal transduction / virulence regulator |
| 12 | PTTG_09156 | PHIG:3617 | *Cryptococcus neoformans* | Protein CPL1 | 29.412 | 4.11E-11 | Signal transduction / virulence regulator |
| 13 | PTTG_12335 | PHIG:5471 | *Puccinia striiformis* | Secreted protein | 27.5 | 8.35E-10 | Secreted protein |
| 14 | PTTG_01817 | PHIG:3617 | *Cryptococcus neoformans* | Protein CPL1 | 27.333 | 2.78E-09 | Signal transduction / virulence regulator |
| 15 | PTTG_09169 | PHIG:3617 | *Cryptococcus neoformans* | Protein CPL1 | 27.742 | 8.86E-09 | Signal transduction / virulence regulator |
| 16 | PTTG_06720 | PHIG:3617 | *Cryptococcus neoformans* | Protein CPL1 | 32.184 | 4.09E-07 | Signal transduction / virulence regulator |

### Identification of Infection-responsive PtEPs

To identify and elucidate putative PtEPs involved during infection, we re-analysed publicly available transcriptomic data from the NCBI Sequence Read Archive (SRA). RNA-seq data from various stages of plant-pathogen interactions were employed to assess the expression profiles of predicted effectors. A subset of PtEP genes demonstrated significant changes in expression in response to infection. The expression of all 273 effector genes was not detected. Among these, 38 and 150 effector genes were identified in expression profiles curated from Singh et al, (2017) and Yadav et al, (2016), respectively **(Figure 4A and 4B)**. Notably, PTTG_05378, followed by PTTG_04313, exhibited significantly higher expression in two independent transcriptomic studies. Additionally, heatmap analysis revealed that several effector genes were differentially expressed across various time points, with distinct patterns observed in early (R0, S0) versus late (R72, S72) infection stages. The expression patterns of these genes underscore their potential involvement in host-pathogen interactions and the facilitation of fungal virulence.

**Figure 4.**
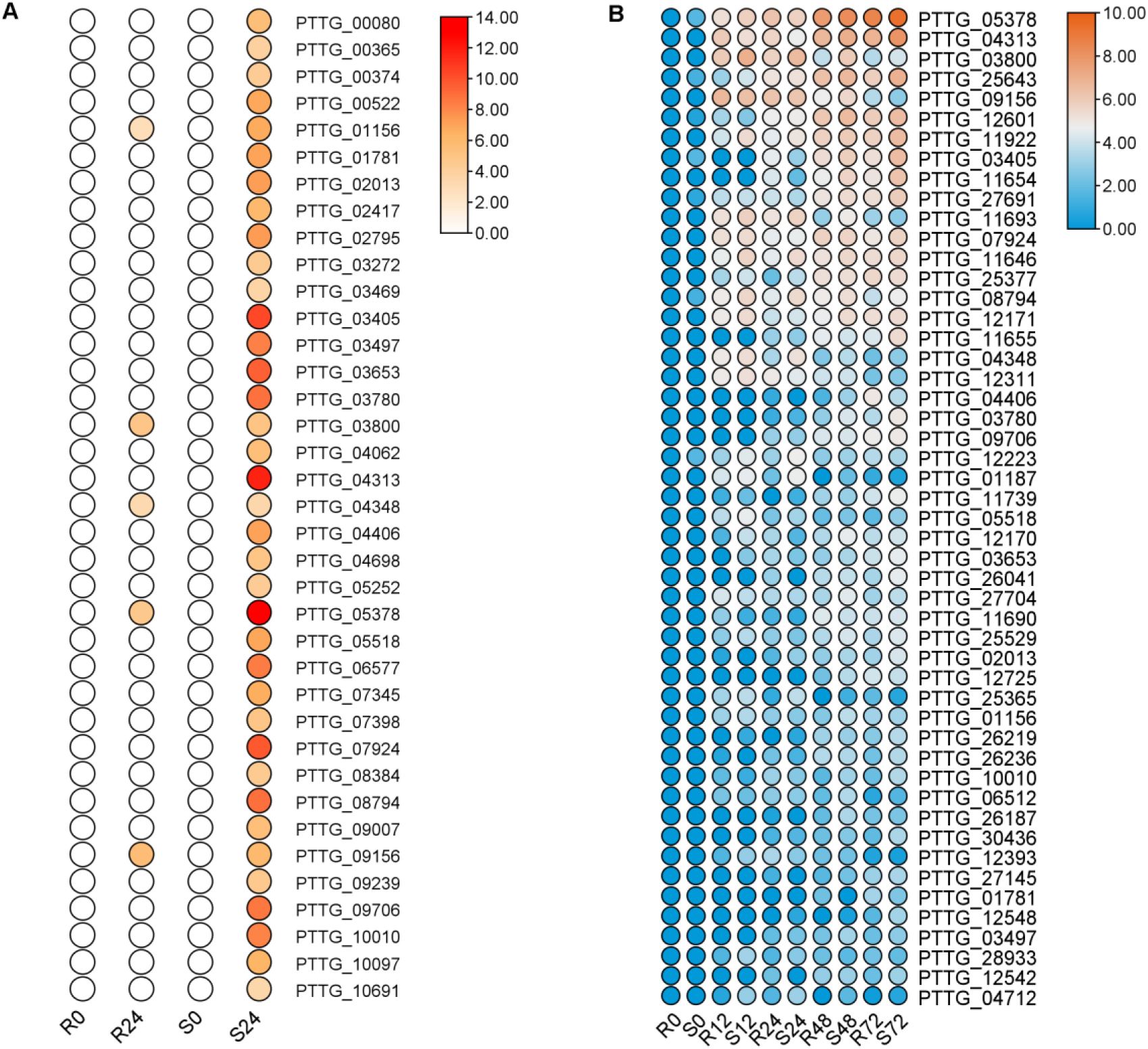
Temporal expression of effector genes in HD2329 + Lr28 resistant and HD2329 susceptible wheat NIL following *P. triticina* infection. (A) Heatmap depicting expression of effector candidates at 0 and 24 hpi (RNA-seq data retrieved from PRJNA182354). (B) Heatmap depicting the expression of *P. triticina* effector genes across multiple infection stages viz 0, 12, 24, 48, 72 hpi (RNA-seq data retrieved from PRJNA328385)

### Gene Structure, and *Cis*-Regulatory analysis of PtEP

Based upon the results so far, genes were selected for further analysis of underlying genomic features. To better understand the diversity inherent in gene structures, the exon-intron organization was analysed utilising full-length effector gene sequences. The genomic configuration of PtEP genes revealed substantial variation in the distribution and length of exons and introns. Notably, some genes contained up to 16 introns, whereas others were entirely devoid of introns **(Figure 5)**. This structural diversity elucidates distinct evolutionary trajectories and characteristics among PtEP genes.

**Figure 5.**
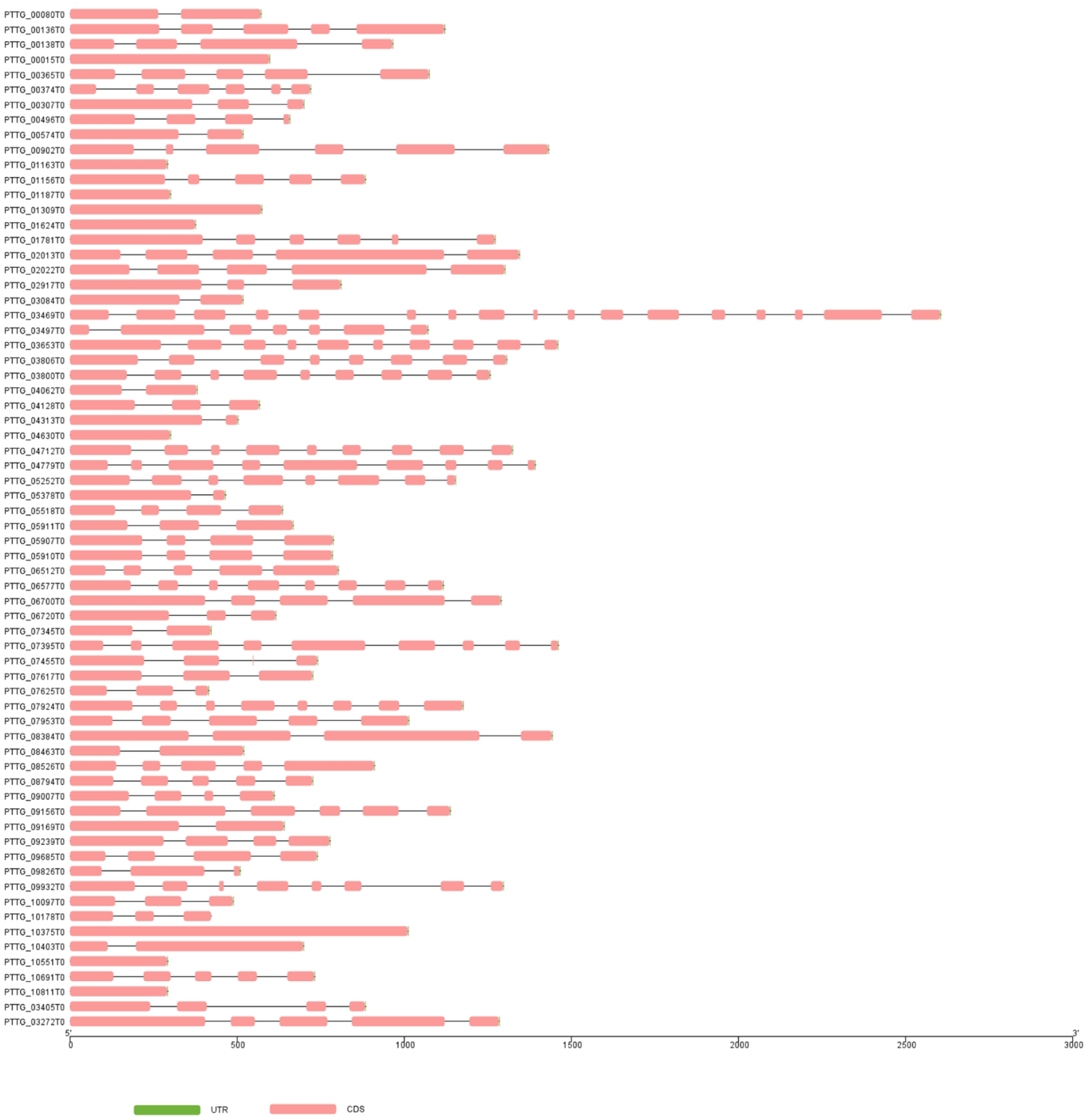
Schematic representation of the genomic architecture of effector genes in *P. triticina*. The exon-intron structure is predicted by Gene Structure Display Server 2.0 (GSDS). The blocks represent coding sequences (pink colour) and UTR region (green colour), while introns are represented by a line. The scale depicts the size of full-length gene along with UTRs.

Furthermore, to elucidate the regulatory mechanisms governing PtEP genes, an analysis of *cis*-regulatory elements within their promoter regions was performed. The distribution of up to 10 motifs was identified using MEME programme within the promoter region **(Figure 6A, Supplementary Figure 4)**. The analysis revealed the presence of 131 cis-acting elements in the PtEP promoter. These regulatory elements can be categorised into four major functional groups: stress and environmental responses, metabolism and nutrient regulation, development and morphogenesis, and cell cycle regulation **(Figure 5B)**. Additionally, transcription factor binding site analysis showed that C6 zinc cluster transcription factors were most abundant, followed by C2H2 zinc finger transcription factors. Other transcription factor families including bZIP, bHLH, Homeobox, and MADS-box proteins were also identified **(Figure 5C)**. These transcription factors are predominantly associated with gene expression related to stress tolerance and play a vital role in cellular and metabolic regulation. Therefore, the enrichment of stress-responsive regulatory elements and key TF-binding sites suggests that these genes are under complex transcriptional control, facilitating dynamic regulation during infection and adaptation to host environments.

**Figure 6.**
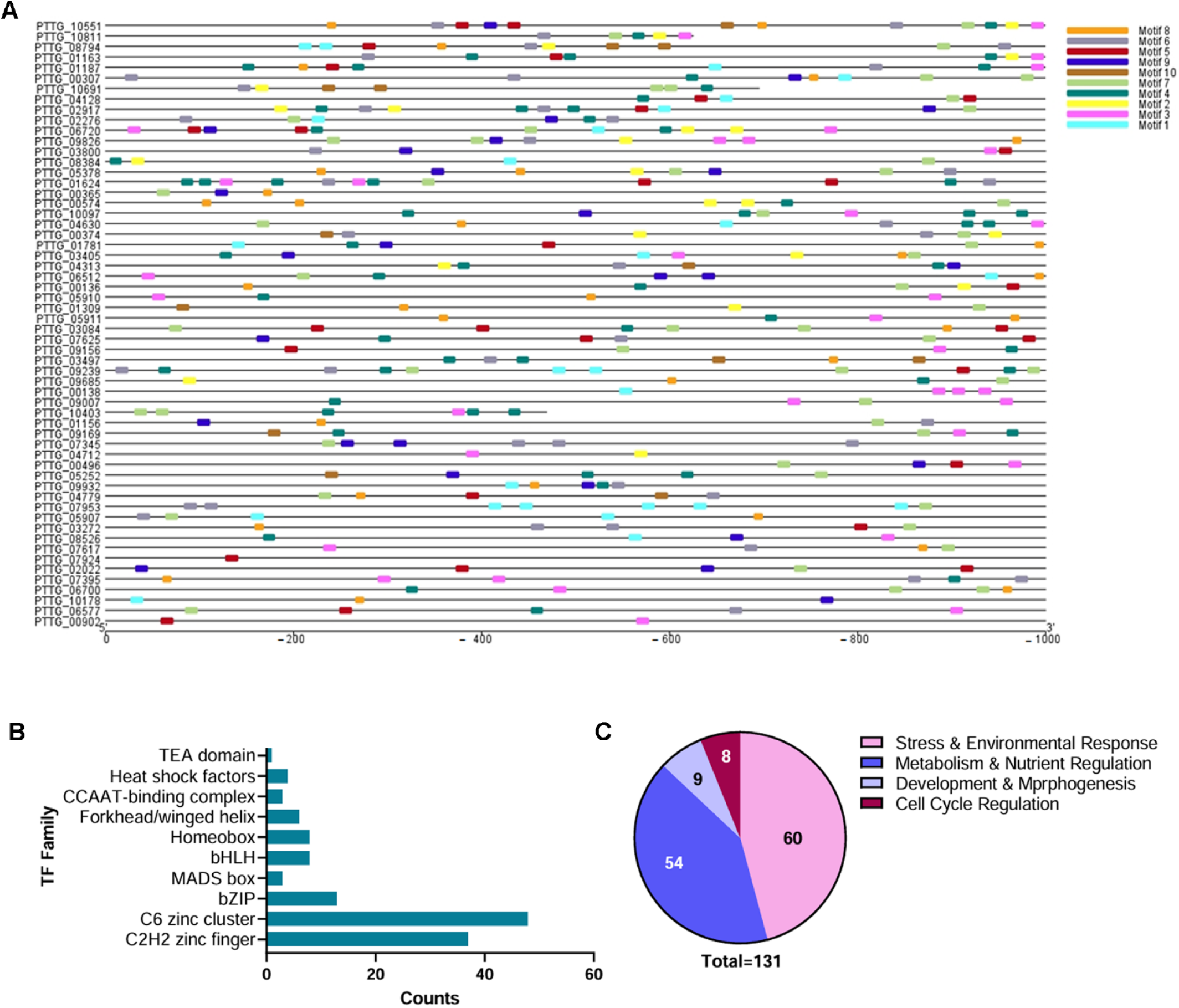
Distribution of *cis*-regulatory elements in the promoter of effector genes. (A) In silico identification of cis-regulatory elements that bind within the promoter region of effector genes. The line depicts promoter length, and colored boxes represent the position of consensus motifs corresponding to transcription factor binding sites. (B) Enrichment in transcription factor families binding to the promoter region. (C) Function governing by transcription factor associated with the identified *cis*-regulatory elements.

### Expression validation and structural characterization of selected Infection-responsive PtEPs

To ensure the accuracy and reproducibility of our transcriptome analysis and to minimise computational bias, we performed a sensitive expression analysis using qRT-PCR. We selected 4 promising disease-responsive candidate effector genes based on the transcriptome data. Their relative expression levels were measured at 0, 12, 24, 72, and 168 hpi, and the fold changes were compared with the RNA-seq data. The qRT-PCR analysis demonstrated expression trends consistent with the transcriptome results. We observed that the transcript levels of the selected genes increased following *P. triticina* infection in comparison to the resistant plant **(Figure 6A)**. Some genes exhibited higher expression levels as the fungal infection progressed **(Figure 6A- ii and iii)**, while others displayed a biphasic pattern of expression **(Figure 6A- i and iv)**.

The extent of expression of these candidate effector genes suggests their potential involvement in the infection process during host-pathogen interactions.

To further investigate their structural features, the mature protein sequences (excluding signal peptide) of the candidate effectors were subjected to structural predictions using the I-TASSER server **(Figure 6B)**. Additionally, we unraveled the underlying information on these effector proteins. Two of the effectors, PTTG_05378 displayed structural resemblance to a hydrolase-like protein, with reliable similarity scores. One effector, PTTG_07924, exhibited strong structural similarity to a Kiwellin protein, and effector PTTG_09156 showed structural similarity to a receptor tyrosine-like kinase **(Table 3)**. Overall, the structural diversity among the modelled effector proteins, as well as their distinct ligand-binding sites, indicates the possibility that they may function through various mechanisms of infection.

**Table 3:**
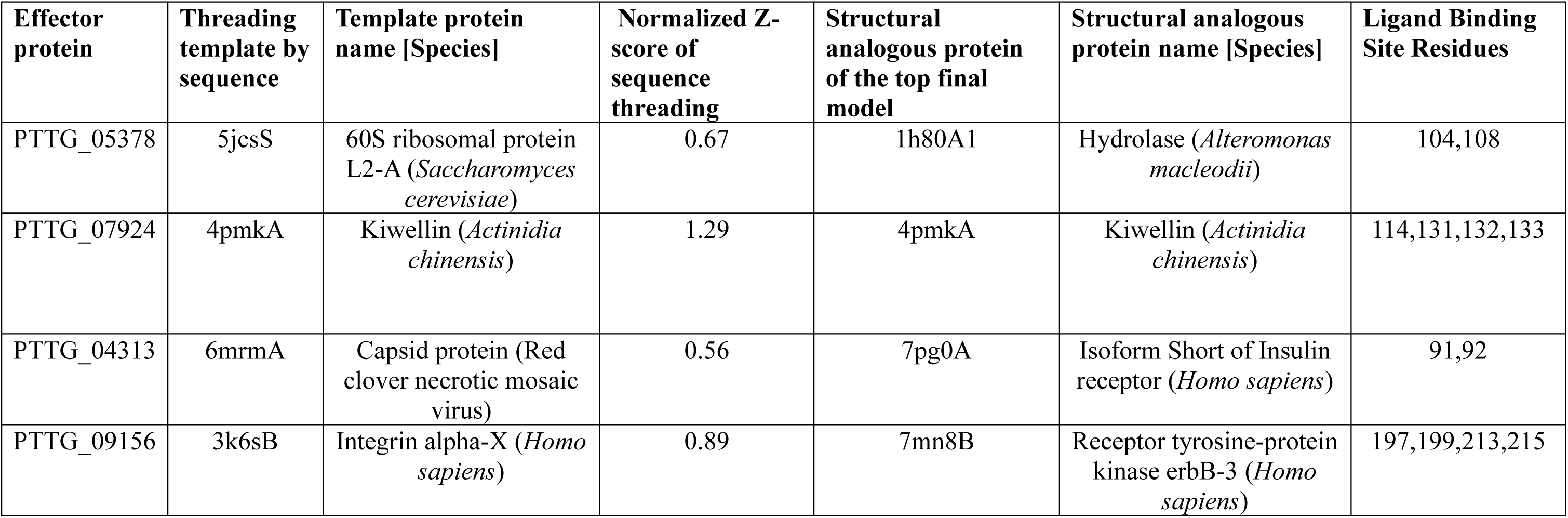
The structurally related proteins in PDB that were selected as template for the prediction of *P. triticina* effector protein structure by I-TASSER server.

## Discussion

Phytopathogenic effectors play an indispensable central role in host invasion, nutrient acquisition, self-protection, and growth (Kamoun, 2006; Win et al., 2012; Rovenich et al., 2014). Fungal effectors, especially those capable of weakening the host’s physical barrier and interfering with the basal immune response, play a vital role in host-pathogen interactions (Jones & Dangl, 2006; Dodds & Rathjen, 2010; Giraldo & Valent, 2013; Lo Presti et al., 2015; Singh et al., 2023). Recent advances in bioinformatics, combined with genomics and transcriptomics, have enabled in-depth comparative analyses of fungal secreted and effector proteins (Sperschneider et al., 2018; Torres et al., 2019). Overall, functional information about *P. triticina* effectors and their role in virulence remains limited (Rampitsch et al. 2015; Figueroa et al. 2016). The majority of *P. triticina* effectors are either uncharacterized or hypothetical. This highlights an exciting opportunity for future research to uncover their mechanisms of fungal virulence.

Effector proteins harbouring conserved sequence features or signature motifs refered as canonical effectors, facilitate their identification and functional prediction. However, due to the rapid evolution resulting in high sequence diversity, many of the effectors are devoid of conserved features or motifs referred as as non-canonical effectors. Prediction of such effectors is solely based on structural or machine learning-based approaches (Stergiopoulos & de Wit, 2009; Lo Presti et al., 2015; Sperschneider et al., 2015, 2018; Win et al., 2012; Petre & Kamoun, 2014). The absence of signature effector-related motifs in certain PtEPs, combined with the limited presence of consensus motifs, highlights that some effector proteins adhere to the classical effector pattern, whereas others are classified as non-canonical or novel effectors lacking well-characterized motif features **(Figure 1)**. Moreover, domain distribution indicates possible functional specialization among effector proteins, which exhibit a broad spectrum of molecular functions **(Figure 2)**. Notably, the absence of conserved domains in PtEPs aligns with the typical characteristics of effector proteins, reflecting their rapid evolution and the lack of well-defined structural features (Stergiopoulos and de Wit 2009).

In the biological process category, the majority of proteins were enriched in areas like “response to other organisms,” “defense response,” “response to hormones,” and “response to abiotic stimuli.” This suggests their important roles in host-pathogen interactions and stress-related responses, highlighting how pathogens adapt and defend themselves. In the molecular functions category, highlighted key terms include “binding,” “protein serine/threonine kinase activity,” and “endopeptidase inhibitor activity”. These functions suggest that effector proteins may interact with host targets, help regulate signalling pathways, and even interfere with host proteolytic processes to facilitate infection (Dodds and Rathjen 2010; Giraldo and Valent 2013). In the cellular component category, most proteins, consistent with the secretory nature of effector proteins, were associated with the “extracellular region” and “extracellular space”. This localization pattern further supports their role as effectors that target host tissues. Furthermore, the activity in genetic information processing and signal transduction pathways indicates that these proteins could help regulate the host cellular machinery during infection. These findings suggest that effector proteins may manipulate host metabolism and signalling pathways for successful infection **(Figure 3A)**. Interestingly, a large number of proteins mapped to KEGG pathways can be attributed to the broad and overlapping nature of pathway annotations. This suggests effector proteins often retain fundamental biochemical functions. Furthermore, KEGG annotations are primarily based on sequence homology, which may result in multiple pathway assignments for individual proteins. Overall, the KEGG analysis highlights the functional diversity of the predicted effector proteins and suggests their involvement in multiple biological processes related to host interaction and pathogenicity **(Figure 3B)**. The CAZyme-containing effectors orchestrate important functions, including host cell wall degradation, modification of carbohydrate structures, and nutrient acquisition during infection (Kubicek et al., 2014; Zhao et al., 2013). The relatively small number of CAZymes-associated proteins compared to KEGG annotations suggests that limited proportion of effector proteins are directly involved in carbohydrate-active mechanism **(Table 1)**. This observation aligns with the concept that only a specific subset of effector proteins functions as carbohydrate-active enzymes, whereas others operate via alternative mechanisms such as signaling interference and immune suppression.

The genomic distribution pattern of *P. triticina* effector genes reveals diverse exon-intron organisation, with some genes harbouring as many as 16 introns, while others remain devoid of introns **(Figure 5)**. An intron-rich gene undergoes complex regulation and may exhibit multiple splice variants, often leading to functional diversity (Reddy et al., 2013; Keren et al., 2010; Zhu et al., 2025). On the contrary, intronless genes, or those with minimal introns, are frequently associated with stress responses and pathogenicity, as they enable faster transcription by requiring little or no splicing (Jeffares et al., 2008; Keeling & Slamovits, 2014), a characteristic of effector proteins. In contrast, genes containing more introns are subject to complex regulation, including alternative splicing, thereby contributing to diversification (Guo et al., 2024). This molecular diversity supports various roles in host-pathogen interactions (Zhu et al., 2025). Furthermore, variation in gene structure underscores the evolutionary flexibility and potential for functional specialization of PtEPs in host-pathogen interaction.

The transcript level expression of selected gene upon following *P. triticina* infection was also documented. Effector genes demonstrate a biphasic pattern of expression (Schneider et al., 2011). The observed increase in transcript levels of selected effector genes at later stages following *P. triticina* infection indicates a significant and sustained fungal virulence response. Conversely, certain genes display a biphasic expression pattern characterised by an initial upregulation, subsequent decline, and a later resurgence, thereby reflecting more complex regulatory mechanisms throughout the infection process **(Figure 7)**. Structural modelling of these effector proteins revealed similarities to a variety of functionally diverse proteins, despite low sequence conservation.

**Figure 7.**
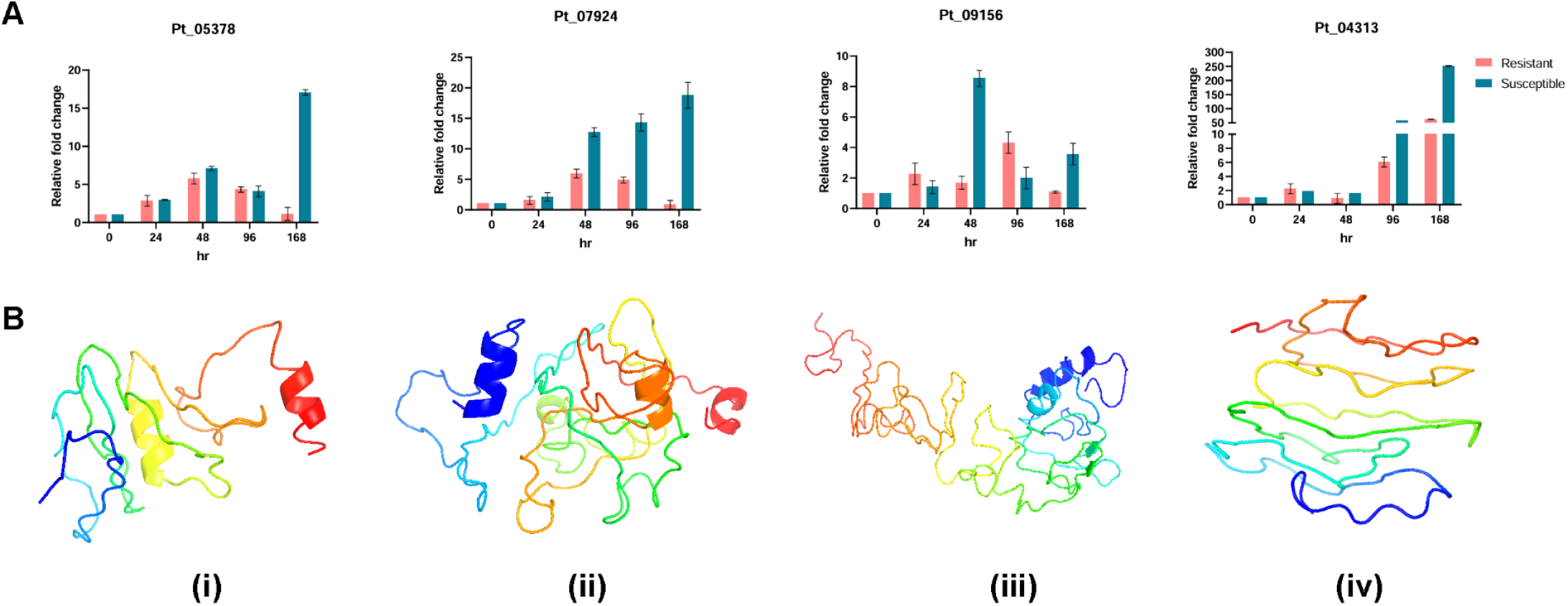
Relative expression and structural analysis of effector candidates. (A) Bar graph showing relative fold change in transcript level of *P. triticina* effector genes at 0, 24, 48, 96, and 168 hpi during leaf rust infection. (B) Structure analysis of candidate effectors whose expression was high in DEGs and expression was validated using qRT-PCR. The three-dimensional structure was obtained through I-TASSER, and the model was displayed through PyMOL.

## Conclusion

In conclusion, we have performed a comprehensive secretome analysis of the P. triticina genome (version GCA_000151525.2) using an *in silico* approach. We predicted a total of 273 effector proteins and catalogued the cytoplasmic and apoplastic effectors of P. triticina. Of these, 17.22% were annotated, while ∼72.22% remain unannotated, indicating a large set of novel effector repertoire. The annotated effectors were assigned roles in metabolism, host interaction, and cellular processes, and limited similarities in PHI-base analysis further support the uniqueness of these effector proteins. A subset of effector proteins was predicted to function as CAZymes, suggesting their potential role in host cell wall modification, a prerequisite for host penetration and pathogenesis. The prioritized effectors showed induced expression during *P. triticina* infection, supporting their involvement in host infection and virulence.

Collectively, these findings provide valuable insights into the characteristics and distinctions of PtEPs, thereby informing future research endeavours into their functional roles. Furthermore, the effector candidates identified in this study, which possess conserved motifs and are upregulated among DEGs upon *P. triticina* infection in the wheat cultivar HD2329, will serve as a valuable resource for future experiments aimed at elucidating the virulence mechanisms of *P. triticina* responsible for leaf rust disease.

## Author contributions

AS and KM curated the project, AS conducted *in silico* analysis. AS performed the wet lab experiments. AS, PK, and HRH assisted with wet-lab experiments. AS and KM analysed the results. AS wrote the first draft. AS, KM, and MK jointly finalized the draft. All authors contributed to the article and approved the submitted version. AS thanks Dr Subhashish Das for helping with the transcriptome analysis and Dr Gaurav Kumar for assistance during manuscript preparation.

## Acknowledgment

The authors would like to acknowledge the Department of Biotechnology (DBT), Government of India, for financial assistance to AS under the grant number DBTRA/2023-24/N/BITS-R/94.

## Supplementary figure

**Supplementary Figure 1:**
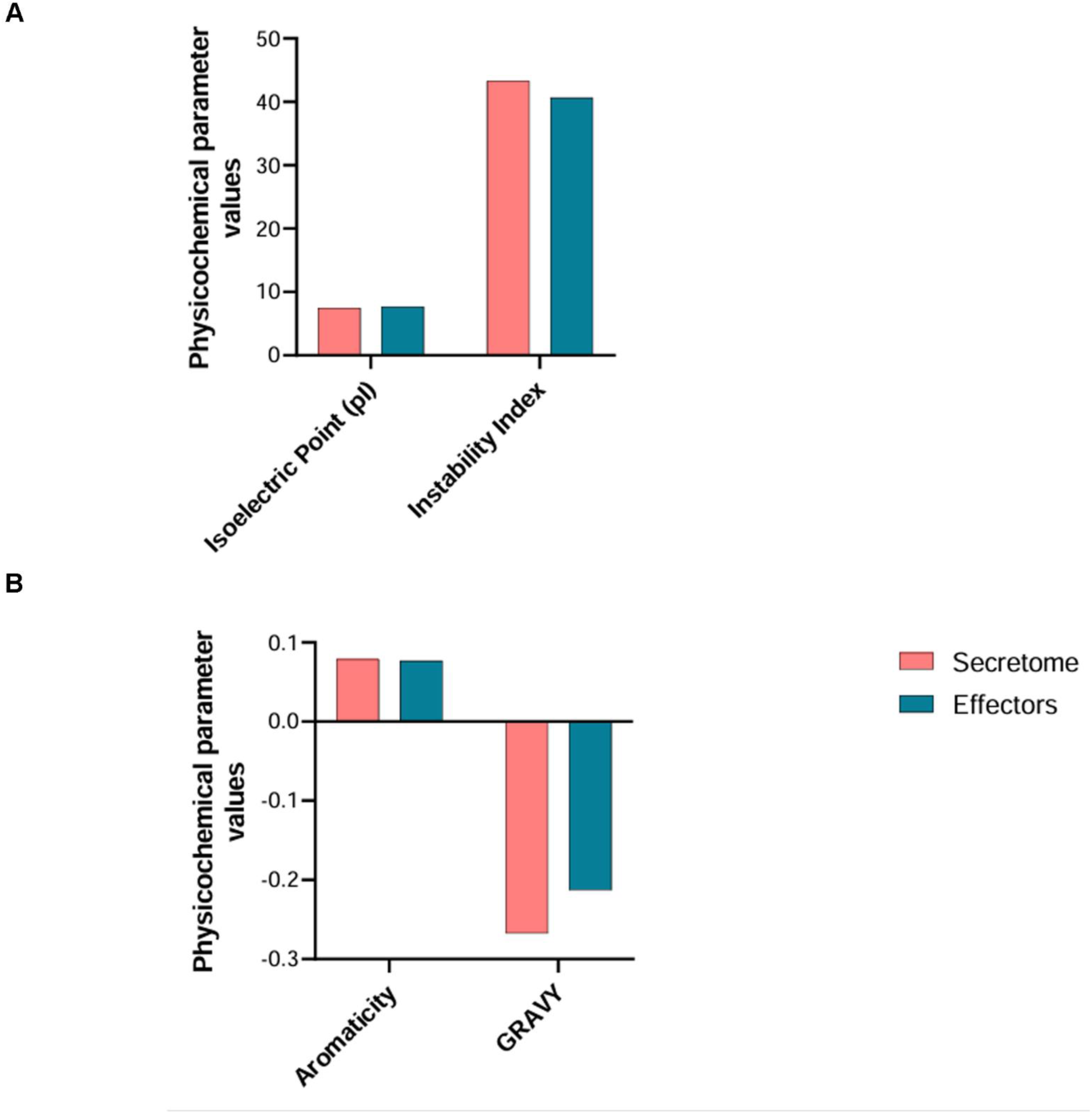
Biochemical property comparison. Bar chart comparing key features (A) Isoelectric point, Instability index, (B) Aromaticity, and GRAVY, between secretory proteins (SP) and effector proteins (EP).

**Supplementary Figure 2:**
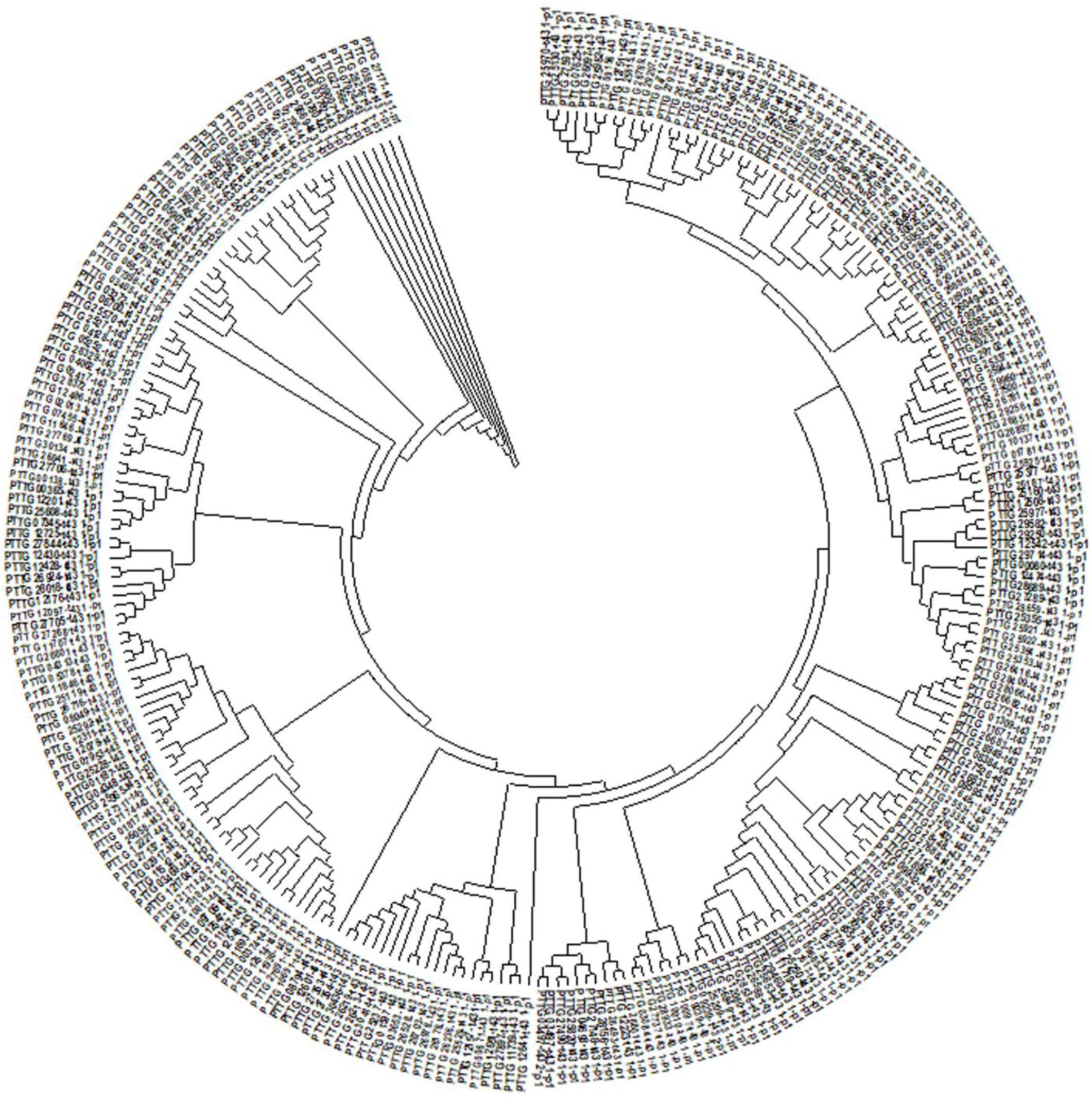
Phylogenetic relationship among PtEPs. A total of 273 proteins were used to construct the phylogenetic tree. The tree was constructed using the Neighbour-joining method, having Bootstrap values from 1,000 iterations

**Supplementary Figure 3:**
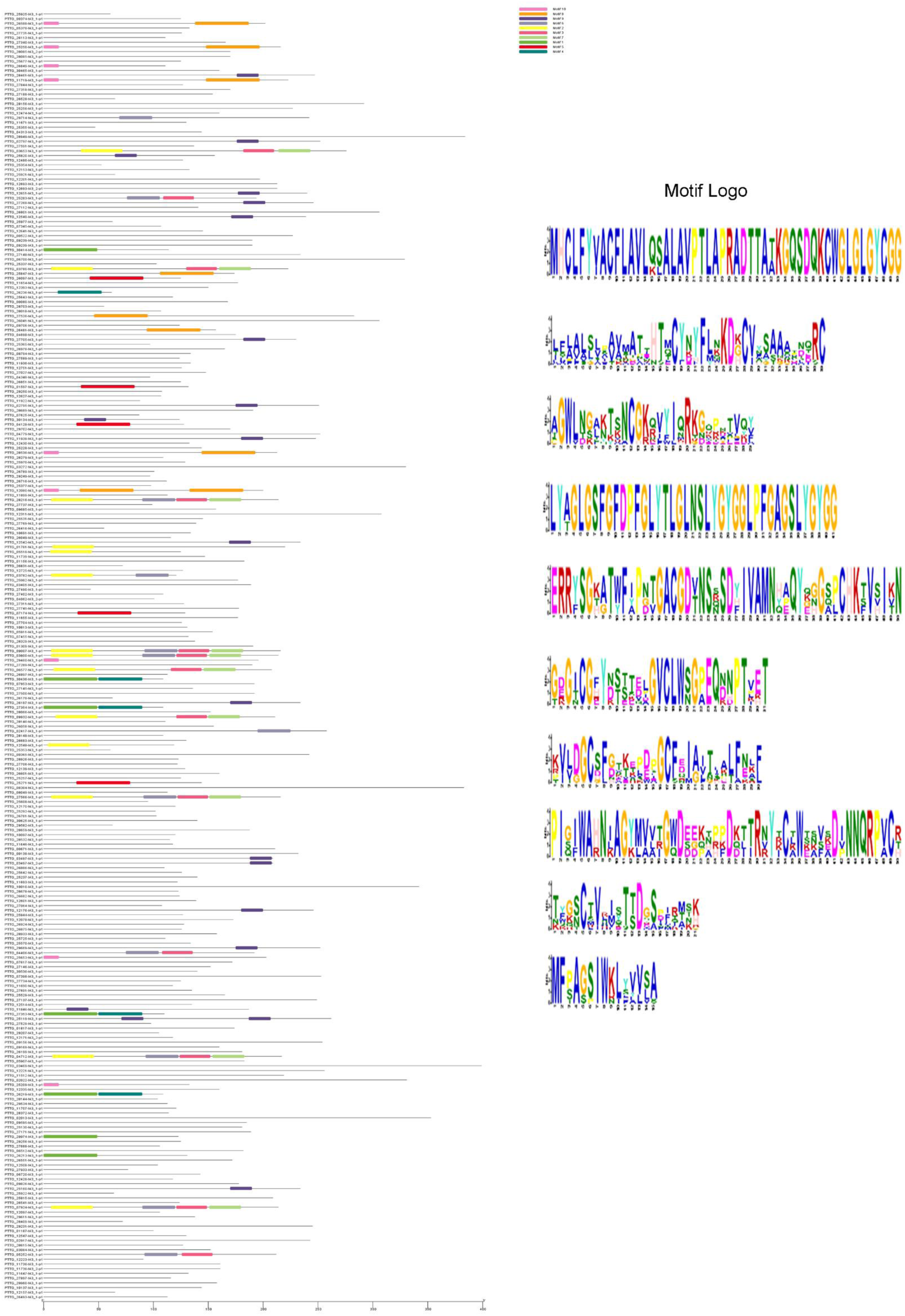
A total of 10 consensus motifs sequences were identified in total 273 effector gene using the MEME programme.

**Supplementary Figure 4:**
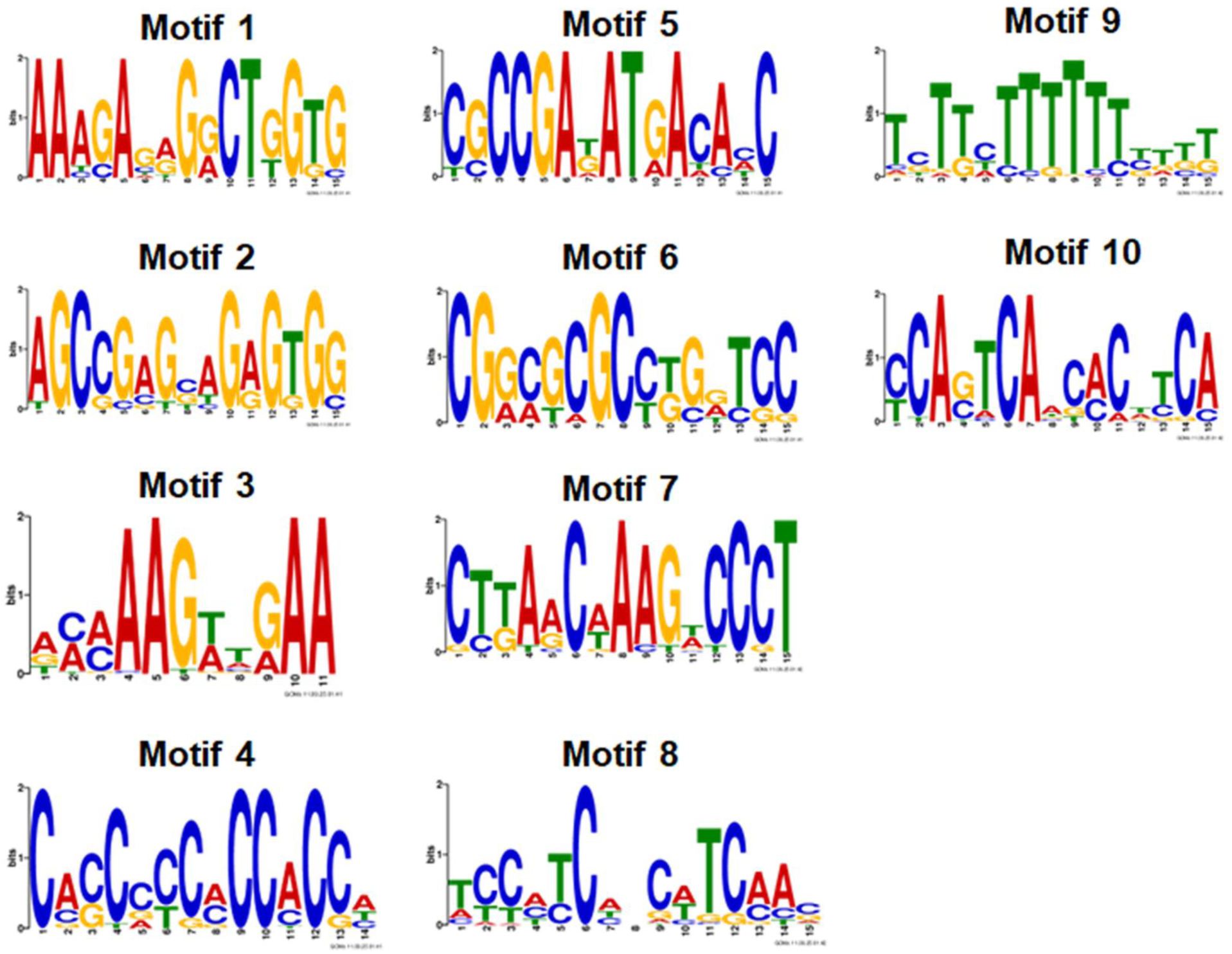
Sequence motif present in the cis-regulatory region, which shows the binding site for various transcription factors.

